# An allosteric agonist activates BK channels by perturbing coupling between Ca^2+^ binding and pore opening

**DOI:** 10.1101/2022.01.28.478270

**Authors:** Guohui Zhang, Xianjin Xu, Zhiguang Jia, Yanyan Geng, Hongwu Liang, Jingyi Shi, Martina Marras, Carlota Abella, Karl L. Magleby, Jonathan R. Silva, Jianhan Chen, Xiaoqin Zou, Jianmin Cui

## Abstract

BK type Ca^2+^-activated K^+^ channels activate in response to both the membrane voltage and intracellular Ca^2+^ with distinct mechanisms. Ca^2+^ binds to the cytosolic domain (CTD) to open the pore across the membrane, but the mechanism that couples Ca^2+^ binding to pore opening is not clear. Here we show that a compound, BC5, identified using *in silico* screening, interacts with BK channels at the interface between the CTD and the transmembrane voltage sensing domain (VSD) and enhances channel activity by specifically affecting the Ca^2+^ dependent mechanism. BC5 activates the channel in the absence of Ca^2+^ binding but Ca^2+^ binding inhibits BC5 effects. Thus, BC5 perturbs the pathway that couples Ca^2+^ binding to pore opening to allosterically affect both, which is supported by atomistic simulations and mutagensis. The results suggest that the CTD-VSD interaction makes a major contribution to the mechanism of Ca^2+^ dependent activation and is an important site for allosteric agonists to modulate BK channel activation.

## Introduction

BK type Ca^2+^ activated K^+^ channels are important in regulating neural excitability^1–3^, neurotransmitter release^4^, and muscle contraction^5, 6^. The aberrant function of BK channels is associated with various human diseases such as neurological disorders^7–9^, hypertension^10^ and abnormal urinary control^11^. The function of BK channels is based on its activation by both elevated intracellular Ca^2+^ and depolarization of the membrane voltage. The opening of BK channels provides a negative feedback mechanism to hyperpolarize the membrane and reduce intracellular Ca^2+^ by deactivating voltage dependent Ca^2+^ channels. On the other hand the Ca^2+^ and voltage dependent activation of BK channels has been an important model system for understanding allosteric mechanisms of ion channel gating^12^. Studies have shown that Ca^2+^ and voltage can activate the channel independently with distinct molecular mechanisms^12–16^. In these activation mechanisms Ca^2+^ or voltage alters the conformation of its sensor, and the conformational change is then coupled to the opening of the activation gate. The understanding of these activation mechanisms will be significant for studying BK channel related physiological processes, developing agents that modulate BK channel activation for therapies, and elucidating fundamental mechanisms of allosteric modulation. However, while the sensors for Ca^2+^ and voltage^13^ and the properties of the activation gate^13, 17^ have been identified, the coupling mechanisms between the sensors and the activation gate in BK channels remain unclear. In this study we address a long-standing issue in the coupling between Ca^2+^ binding and pore opening.

BK channels are formed by four Slo1 subunits with structural domains that are associated with distinct functions ^18–20^ (Fig 1a). In the membrane, a central pore is formed by the transmembrane segments S5-S6 from all four subunits, and the voltage sensor domains (VSDs) from the four subunits, comprising of the transmembrane segments S1-S4, surround the pore. Slo1 also contains an additional transmembrane segment S0. The cytosolic domain (CTD) of each Slo1 subunit harbors two Ca^2+^ binding sites, and the four CTDs form a ring-like structure, known as the gating ring. For Ca^2+^ binding at the CTD and the VSD activation to open the pore, interactions among these structural domains are essential.

**Figure 1.**
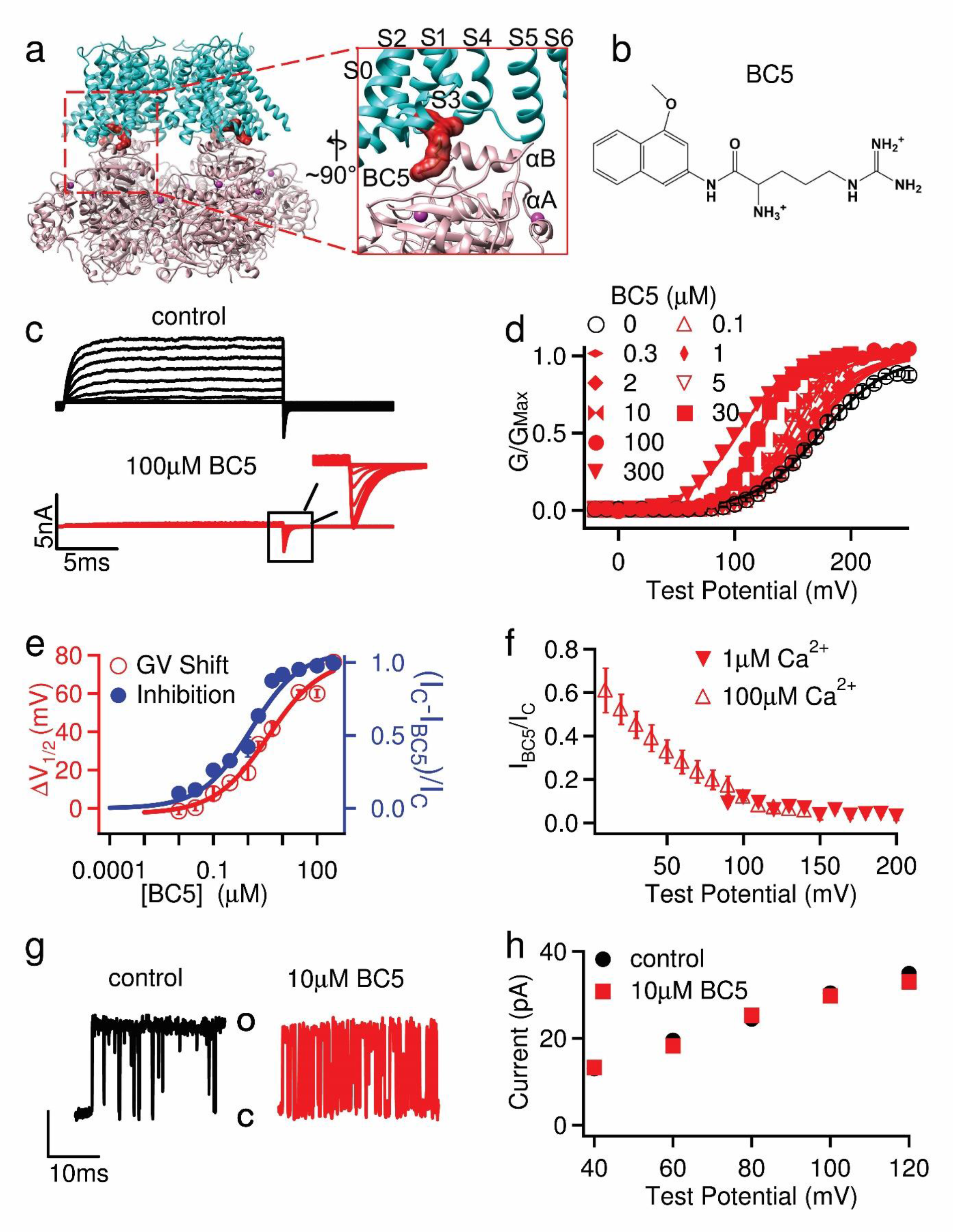
BC5 effects on BK channels. **a,** *hSol1* Channel cryo-EM structure (PDB: 6v38) docked with BC5. The membrane spanning region (S0-S6) from one subunit and the N-terminus of the cytosolic domain (CTD) from a neighboring subunit are amplified in the inset. **b,** BC5 chemical structure. **c,** Current traces of BK channels (control, black) and with BC5 (100 µM, red) at various testing voltages from -30mV to 250mV with 20mV increment. The potential before and after testing pulses was -80 mV. Inset: tail currents after the testing pulses. **d,** GV relationship for WT BK channels at various BC5 concentration. Solid lines are fits to the Boltzmann relation. **e,** The shifts of GV relationships (red, V1/2: voltage where the GV relation is half maximum) and outward macroscopic current inhibition (blue, IC: control current, IBC5: current in BC5) depend on BC5 concentration, with EC50 of 2.5 µM and IC50 of 1.2 µM respectively. **f,** Inhibition of the outward macroscopic current by 100 µM BC5 versus voltage in the presence of either 1µM (filled) or 100 µM (hollow) free Ca^2+^. **g,** Representative single-channel current recordings at 100 mV and in 100 µM free Ca^2+^ as control (left and black) and with 10 µM BC5 (right and red) in an inside out patch in symmetrical 160 mM KCl. O: open; C: closed. **h,** Voltage dependence of single-channel current amplitudes (i-V). The data points represent the mean ± SEM, n ≥ 3 for all figures unless specified otherwise.

What interactions among the structural domains are important for Ca^2+^ dependent activation have been intensively studied but still remain unclear. In the Slo1 subunit the CTD is covalently connected to the S6 by a peptide of 15 amino acid residues, known as the C-linker. A previous study showed that deletion or insertion of residues in the C-linker altered Ca^2+^ and voltage dependent activation in correlation with the C-linker length, which suggested that Ca^2+^ binding to the CTD may open the channel by tugging the C-linker ^21^. On the other hand, the CTD also makes a non-covalent interaction with the VSD of a neighboring subunit. This was first revealed by the discovery that the residues from both the CTD and VSD of the neighboring subunits form the Mg^2+^ binding site for activating the channel^22^, and then was more clearly shown by the structures of the BK channel^18–20^. It was hypothesized that the CTD-VSD interaction may contribute to the coupling between Ca^2+^ binding and pore opening^23^, and the hypothesis seemed particularly strong because the comparison between the structure of the BK channel with Ca^2+^/Mg^2+^ bound and that of metal-free showed a large Ca^2+^/Mg^2+^ dependent change in the interface between CTD and VSD^18–20^. This hypothesis was consistent with an earlier finding that the part of the CTD located close to the VSD is important in determining the different Ca^2+^ sensitivities between different BK channel homologs^24^. Furthermore, mutations in the CTD-VSD interface also altered Ca^2+^ sensitivity^25, 26^. However, these mutations either had small effects on Ca^2+^ dependent activation^26^ or, besides reducing Ca^2+^ dependent activation, also altered voltage dependent activation and resulted in an increase of the intrinsic open probability of the channel^25^, suggesting that the mutations may have caused structural changes that non-specifically affected the overall function of the channel.

In this study we have used *in silico* screening to identify a compound that binds to a site at the CTD-VSD interface and enhances BK channel activation. We found that the compound activates the channel by perturbing the pathway for coupling between Ca^2+^ binding and pore opening. These results support the idea that the CTD-VSD interaction makes a major contribution to the coupling between Ca^2+^ binding and pore opening, and the CTD-VSD interface is an important site for allosteric agonists to enhance Ca^2+^ dependent activation of BK channels.

## Results

In order to probe if the VSD-CTD interaction of BK channels is important for Ca^2+^ dependent activation, we initiated a search for compounds that may interfere with BK channel Ca^2+^ dependent activation *in silico*. For this, we docked a subset of the Available Chemical Database (ACD, Molecular Design Ltd.) of about 4×10^4^ compounds, in which each compound has a formal charge of either 1 or 2, to a site near the VSD-CTD interface (Fig 1a). Patch-clamp recordings of BK channels expressed in *Xenopus* oocytes with and without the candidate compounds from the *in silico* screening showed that BC5 (Fig 1b) activated the channel as measured from the inward tail currents at a voltage (-80 mV) more negative than the equilibrium potential of the K^+^ (0 mV) (Fig 1c, d, sFig 1a). BC5 shifted the voltage dependent activation (GV relation) of BK channels to more negative voltages (Fig 1d), with the concentration at the half-maximum GV shift (EC50) of 2.5 µM (Fig 1e). On the other hand, BC5, applied in the cytosolic side of the membrane patch, also inhibited the outward BK currents at positive voltages (Fig 1c, sFig 1b).

We found that the BC5 activation and inhibition of BK channels were independent molecular processes. BC5 inhibited the channel at a lower concentration than activation, with the half-maximum inhibition concentration (IC50) of 1.2 µM (Fig 1e, sFig 1b). The inhibition was dependent on voltage, with a weaker inhibition at less positive voltages (Fig 1f) and no inhibition at a negative voltage (Fig 1c). These characteristics of inhibition are consistent with the mechanism that the positively charged BC5 acts as a pore blocker from the cytosolic side, and can be flushed out by negative voltages and inward K^+^ currents, which is reminiscent of Ca^2+^ and Mg^2+^ block of the channel^27, 28^. Single channel recordings showed that BC5 caused brief closures during channel opening but without altering the single channel conductance (Fig 1g, h), consistent with the mechanism that BC5 acts as a pore blocker with fast on and off rates. In addition, unlike BC5 activation, BC5 inhibition of the channel showed little dependence on Ca^2+^ (Fig 1f) or Mg^2+^ (sFig2a, b), and the mutations that reduced BC5 activation had little effect on BC5 inhibition (Fig 1e, sFig 3a,b). These results suggest that BC5 inhibition of the channel is an off-target effect, such that, in addition to interacting with our targeted docking site for channel activation (Fig 1a), BC5 may interact with the pore (sFig3c) to block the outward K^+^ currents.

Our targeted docking site of BC5 in BK channels is located close to the Mg^2+^ binding site for channel activation, which is formed by residues from both the VSD and CTD^22, 29^ (Fig 2a). As shown in Fig 2a, d, residue E399 is also shared by both the Mg^2+^ and BC5 binding sites. If BC5 interacts with the targeted docking site to activate the channel, we expected that an electrostatic repulsion between Mg^2+^ and the charges in BC5 and a direct competition between Mg^2+^ and BC5 for binding would reduce BC5 activation. Consistent with this idea, we found that BC5 at the same concentration induced less shift in GV relations in the presence of 10 mM Mg^2+^ (Fig 2b, c, sFig 2a). Likewise, a charge reversal mutation of the Mg^2+^ binding residue, D99R^19^, also reduced BC5 activation of the channel (Fig 2c), supporting that BC5 binds in the vicinity of the Mg^2+^ binding site.

**Figure 2.**
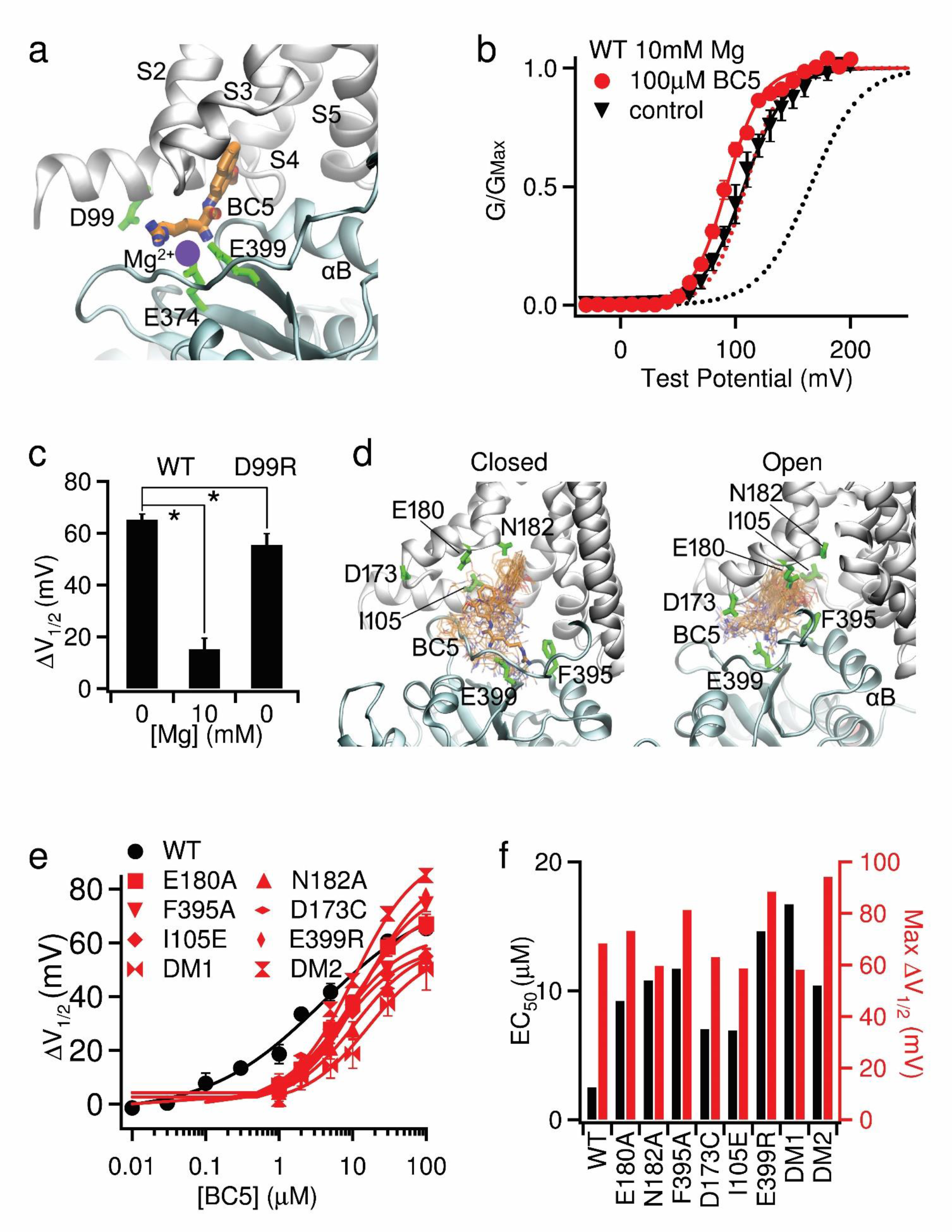
BC5 interactions with the targeted site at the CTD-VSD interface. **a,** The cryo-EM structure of *hSlo1* (PDB: 6v38) with BC5 docked to its targeted site. The VSD and CTD are colored in silver and cyan, respectively. Mg^2+^ binding residues (D99, E374, E399) are represented as green sticks. The Mg^2+^ position is highlighted as the blue circle. **b,** GV relationship of BK channels and Boltzmann fits (solid lines) in 10 mM Mg^2+^. V1/2 and slope factor (mV) for control: 107.4 ± 3.8 and 18.3 ± 3.4; and 100 µM BC5: 92.2 ± 3.7 and 15.0 ± 3.3. For comparison, the dashed lines represent the GV relationships of the control (black) and 100 µM BC5 (red) in the absence of Mg^2+^, taken from Fig 1d. **c.** GV shifts caused by 100 µM BC5. For WT channels the GV shifts in 0 and 10 mM Mg^2+^ are plotted, and for the D99R mutant channels, GV was measured in 0 Mg^2+^. *: p<0.05, unpaired Student’s t-test. **d**. Dynamic interactions of BC5 in the binding pocket of metal-free (left) and Ca^2+^-bound (right) BK channels. Snapshots of BC5 (thin sticks) were extracted from the atomisitic simulation trajectories every 10 ns and superimposed onto the last frame of *sim 1* and *7* (Table S1). **e,** GV shifts of mutant BK channels in response to BC5. Each data point was averaged from recordings in 3-9 patches. The WT curve is taken from Fig 1E. **f,** Maximal G-V shift and EC50 of BC5 for mutations and the WT. DM1: double mutation E180A/N182A; DM2: I105E/E399R.

To further characterize BC5 interactions with the targeted docking site, 200 ns molecular dynamics (MD) simulations of the open and closed states of BK channel with and without docked BC5 were performed in explicit membrane and water (see Methods) (sTable 1). The results showed multiple binding modes between BC5 and the channel that interchanged dynamically with time, and in the process BC5 interacted with different residues (Fig. 2d). As summarized in sTable 2, the hydrophobic naphthalenyl group of BC5 was buried in contact with the intracellular side of S0, S1 S3 and S4 as well as the residue F395 from αB of CTD. The charged arginine part of BC5 interacted with the negatively charged residue E399 (Fig 2d). The binding poses of BC5 are more narrowly defined in the open conformation, suggesting that BC5 binding is stronger and may favor the open state of BK channels. We mutated the BC5 interacting residues individually or in combination to alanine or other amino acids that changed the charge of the residue, and measured dose responses of the mutant channel activation to BC5 (Fig 2e). These mutant channels showed increased EC50 (Fig 2f), suggesting that the mutations reduced BC5 binding by interrupting the interactions between the residue and BC5. These results support that BC5 enhances BK channel activation by binding to our targeted docking site.

BK channels are activated by voltage and Ca^2+^ with distinctive mechanisms^7, 14^. Does BC5 shift the GV relation of BK channels by modulating the mechanism of voltage or Ca^2+^ dependent activation? To address this question, we first examined if BC5 modulated VSD activation in BK channels by measuring gating currents with and without BC5 (Fig 3a). Unlike the GV relation, the voltage dependence of gating charge movement (QV) was not shifted to more negative voltages (Fig 3a), suggesting that BC5 did not activate the channel by altering the voltage dependent mechanism. Further supporting this conclusion, measurements of intrinsic openings of the channel at negative voltages (<-100 mV) where the voltage sensors were at the resting state^15, 30^ showed that BC5 enhanced open probability (Fig 3b). Mg^2+^ activates BK channels by interacting with the voltage sensor when the VSD is at the activated conformation^31–33^, which results in a reduction of the off-gating current amplitude at the end of the depolarizing voltage pulse^34^. The Mg^2+^-VSD interaction also shifts GV relation to more negative voltages. However, Mg^2+^ cannot alter open probabilities at negative voltages when the VSD is not activated^34^, differing from the BC5 effect (Fig 3b). Unlike Mg^2+^, BC5 also had no effect on the amplitude of off-gating currents (Fig 3a). Therefore, BC5 did not alter voltage dependent activation indirectly as well by any mechanism similar to that of Mg^2+^. On the other hand, Ca^2+^ binding also shifts the GV relation to more negative voltages and enhances open probability at negative voltages^12, 35^. Interestingly, the amount of GV shift and the enhancement of open probability at negative voltages caused by 100 µM BC5 were comparable to those caused by 1 µM Ca^2+ 14, 35^, suggesting that BC5 activated the channel by modulating the Ca^2+^ dependent mechanism.

**Figure 3.**
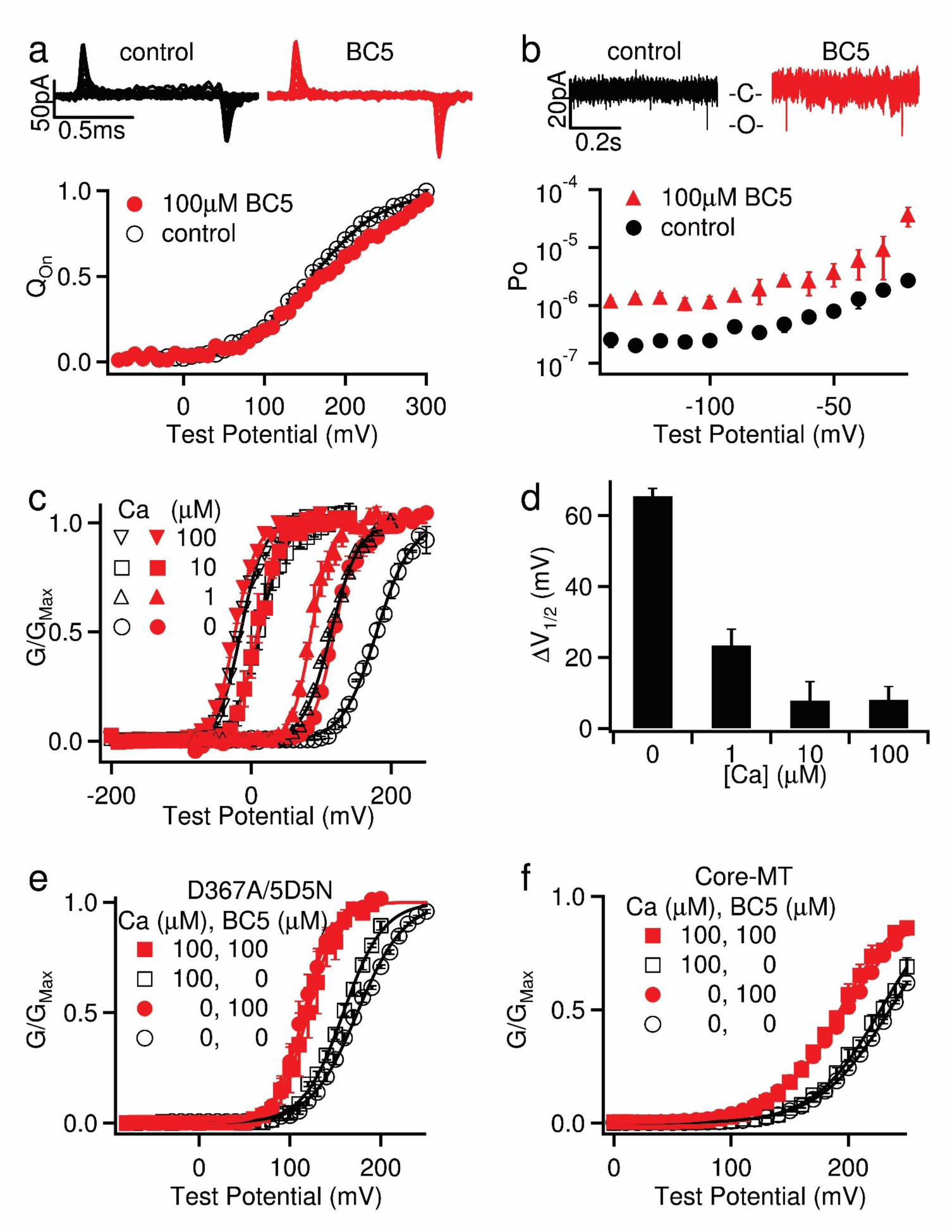
BC5 perturbs Ca^2+^ dependent activation. **a,** Top, gating currents in control and 100 µM BC5. Voltage pulses were from -80 to 300 mV with 20 mV increments. Bottom, Normalized gating charge-voltage (QV) relation of on-gating currents. The smooth lines are fits to the Boltzmann function with V1/2 and slope factor of 169.3 ± 5.0 mV and 49.8 ± 4.6 mV for control, and 185.1 ± 6.5 mV and 56.6 ± 6.3 mV for BC5. **b,** Top, current traces of a patch containing hundreds of BK channels at −140 mV in control and 100 µM BC5. Only brief unitary channel openings were seen at the negative voltage. O: open; C: closed. Bottom, Open probability (Po) at negative voltages. **c,** GV relations in various [Ca^2+^]i (black hollow symbols) and in addition of 100 µM BC5 (red filled symbols). Black solid lines are Boltzmann fits without BC5. V1/2 and slope factor (mV): 175.0 ± 3.4 and 22.1 ± 3.0 (0 µM [Ca^2+^]i), 114.1 ± 3.8 and 19.9 ± 3.4 (1 µM [Ca^2+^]i), 11.3 ± 4.4 and 18.4 ± 3.9 (10 µM [Ca^2+^]i), and -15.1 ± 3.8 and 16.1 ± 3.3 (100 µM [Ca^2+^]i). Red solid lines are Boltzmann fits with BC5. V1/2 and slope factor (mV): 108.7 ± 3.0 and 15.0 ± 2.7 (0 µM [Ca^2+^]i), 85.9 ± 3.8 and 13.7 ± 3.3 (1 µM [Ca^2+^]i), 8.8 ± 3.3 and 15.9 ± 2.9 (10 µM [Ca^2+^]i), and -23.1 ± 3.3 and 15.6 ± 2.5 (100 µM [Ca^2+^]i). **d,** GV shifts in response to 100 µM BC5 at different [Ca^2+^]i. **e,** BC5 effects on the D367A5D5N mutant channel in 0 and 100 µM [Ca^2+^]i . D367A and 5D5N (D897-901N) ablated the two Ca^2+^ binding sites in each Slo1 subunit, respectively. GV relations were fit with the Boltzmann function (solid lines). V1/2 and slope factor (mV): 174.2 ± 4.2 and 25.7 ± 3.8 for 0 [Ca^2+^]i and 0 BC5; 115.4 ± 3.0 and 15.8 ± 2.6 for 0 [Ca^2+^]i and 100 µM BC5; 158.8 ± 3.1 and 22.1 ± 3.0 for 100 µM [Ca^2+^]i and 0 BC5; and 119. 9 ± 3.1 and 16.3 ± 2.7 for 100 µM [Ca^2+^]i and 100 µM BC5. **f,** BC5 effects on the Core-MT BK channel in 0 and 100 µM [Ca^2+^]i. . GV relations were fit with the Boltzmann function (solid lines). V1/2 and slope factor (mV): 235.5 ± 3.1 and 30.4 ± 3.2 for 0 [Ca^2+^]i and 0 BC5; 198.1 ± 3.0 and 31.7 ± 3.0 for 0 [Ca^2+^]i and 100 µM BC5; 227.5 ± 3.8 and 28.4 ± 4.0 for 100 µM [Ca^2+^]i and 0 BC5; and 192.9 ± 3.3 and 27.8 ± 3.1 for 100 µM [Ca^2+^]i and 100 µM BC5.

To further examine the BC5 perturbation of the mechanism of Ca^2+^ dependent activation, we measured BC5 activation of BK channels at various Ca^2+^ concentrations. We found that Ca^2+^ reduced BC5 activation and the reduction was dependent on Ca^2+^ concentration. At 100 µM Ca^2+^, which saturated Ca^2+^ dependent activation^11^, BC5 no longer activated the channel (Fig 3c, d, sFig 4). The effect of Ca^2+^ on BC5 activation is specifically associated with Ca^2+^ dependent activation of the channel, because mutations that abolished Ca^2+^ binding sites in the CTD for channel activation^36^ also abolished Ca^2+^ effects on BC5 activation of the channel (Fig 3e, sFig 5). We studied a truncated BK channel in which the entire CTD was replaced with a short exogenous peptide, known as the Core-MT^37^. The Core-MT channel was no longer activated by Ca^2+^ because the deletion of all Ca^2+^ binding sites. However, it was still activated by BC5, and 100 µM Ca^2+^ had no effect on BC5 activation (Fig 3f, sFig 6a-c). These results suggest that Ca^2+^ competed with BC5 for activating BK channels.

**Figure 4.**
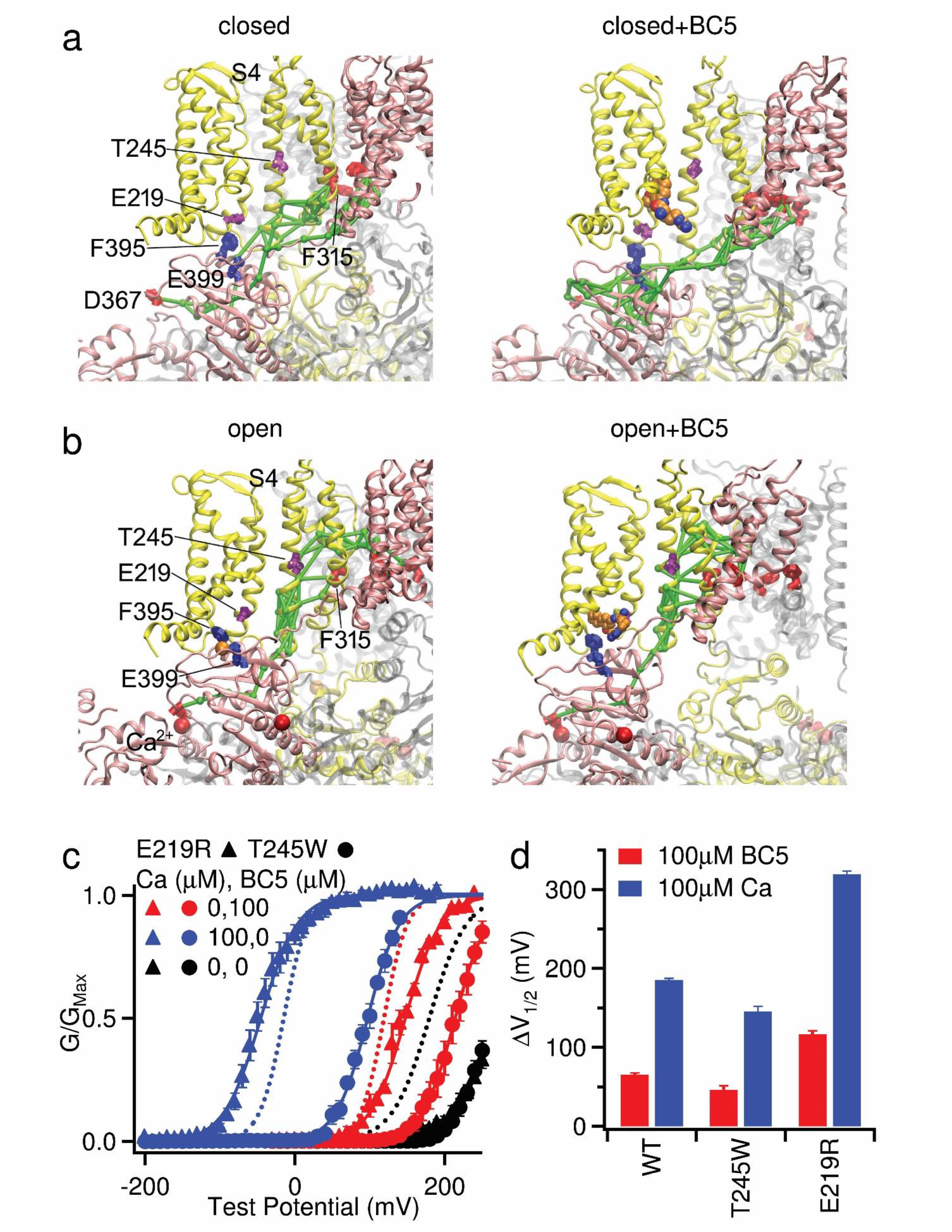
BC5 interactions with the allosteric pathway for coupling Ca^2+^ binding to pore opening. **a,b,** Optimal and suboptimal pathways coupling Ca^2+^ binging site D367 and pore lining residue F315. The two neighboring subunits forming the VSD-CTD contacts are colored in pink and yellow, respectively. The pathways are represent in green sticks. **c,** GV relationship and Boltzmann fits (solid lines) of the mutant channels E219R (results are shown with triangles) and T245W (results are shown with circles). V1/2 and slope factor (mV): for E219R, 263.7 ± 8.7 and 30.4 ± 6.8 at 0 [Ca^2+^]i and 0 BC5 (black triangles); 147.4 ± 4.7 and 23.6 ± 4.1 at 0 [Ca^2+^]i and 100 µM BC5 (red triangles); and -51.5 ± 4.2 mV and 24.7 ± 3.5 at 100 µM [Ca^2+^]i and 0 BC5 (blue triangles); for T245W, 260.9 ± 6.8 and 21.1 ± 5.3 at 0 [Ca^2+^]i and 0 BC5 (black circle); 213.8 ± 2.9 and 22.1 ± 2.8 W at 0 [Ca^2+^]i and 100 µM BC5 (red circle); and 106.0 ± 3.3 and 21.2 ± 3.1 at 100 µM [Ca^2+^]i and 0 BC5 (blue circle). Dashed lines are taken from Fig. 3C for WT at 0 [Ca^2+^]i and 0 BC5 (black); at 0 [Ca^2+^]i and 100 µM BC5 (red); and at 100 µM [Ca^2+^]i and 0 BC5 (blue). **d,** GV shift caused by 100 µM [Ca^2+^]i (blue) and 100 µM BC5 (red).

Since Ca^2+^ binding sites in the CTD are not spatially close to the BC5 binding site at the VSD- CTD interface (Fig 1a), and further, BC5 activated the mutant channels in which Ca^2+^ binding sites or the entire CTD were eliminated (Fig 3e, f), BC5 likely does not directly interact with Ca^2+^ binding, but rather the competition between Ca^2+^ and BC5 is allosteric, i.e., Ca^2+^ binding to the channel may alter the conformation at the BC5 binding site. For BC5 to open the activation gate independently of voltage on one hand, and compete with Ca^2+^ binding on the other hand, it may interact with the molecular pathway that couples Ca^2+^ binding to channel opening. That is, BC5 perturbs the pathway to promote pore opening, which can be overwhelmed by a stronger perturbation at high Ca^2+^ concentrations due to Ca^2+^ binding. To further explore this idea, we performed network analysis (Fig 4) and information flow analysis (sFig 7) to examine the coupling pathway from the Ca^2+^ binding residue D367 to the pore lining F315. For this, network were constructed where each node represented a single residue (or BC5) and the lengths of edges captured the dynamic correlation calculated from the MD trajectories (see Methods). Nodes that are more strongly coupled will have shorter connecting edges. We then calculate a set of “optimal” and “suboptimal” paths between the selected residues, which are the shortest paths and thus the most strongly coupled. ^38^ The top 20 paths with the strongest coupling strengths are shown in Fig 4a-d (green sticks) to illustrate the allosteric coupling pathways connecting Ca^2+^ binding site (D367) to the pore residues (F315) in both the same subunit and a neighbor subunit. The Ca^2+^ binding site is in the CTD, while the activation gate is located in the membrane spanning domain. In both the open and closed conformations, the allosteric pathway crossed these domains via two major branches, with one via the VSD-CTD interface from αA/αB in the CTD to the S4-S5 linker/S6 in the VSD, and the other via the covalent C-linker to S6. To complement the pathway analysis, we analyze the information flow^39, 40^ between D367 (“source”) and F315 (“sink”), which quantifies the contributions of all residues to the dynamic coupling. The results show that the information flow through VSD-CTD interface is higher or similar to that through the C-linker/S6 connection (sFig.7), suggesting that these structural elements are critical for the coupling between Ca^2+^ binding and pore opening in addition to the C-linker/S6 pathway^21^. Importantly, the network analysis shows that BC5 lies adjacent to the key allosteric pathways at the VSD-CTD interface, interacting with residues F395 and E399, which are part of both the BC5 binding site and the allosteric pathway in the closed+BC5 structure (Fig 4a). In addition, in other closed and open structures F395 and E399 are in the vicinity of the allosteric pathway within a distance < 6 Å (Fig 4a, b; sTable 2). These results suggest that BC5 binding likely perturbs the allosteric pathway to interact with both Ca^2+^ binding and pore opening.

We performed a mutational scanning of selected residues on or in the vicinity of the allosteric pathway and found that the mutation T245W reduced, while E219R enhanced the sensitivity of the channel to Ca^2+^ for activation^41^ (Fig 4e, f). T245 is part of the allosteric pathway, while E219 is in the vicinity of the allosteric pathway in the open state with or without BC5 (Fig 4b). E219 and T245 are in S4 and S5, respectively, and downstream of BC5 in the allosteric pathway, being closer to the activation gate (Fig 4a-d). If BC5 perturbs the allosteric pathway to activate the channel as shown above (Fig 4a-d; sFig 7) we expected that these two mutations would alter BC5 activation as well. Like the effects on Ca^2+^ dependent activation, T245W reduced BC5 activation of the channel such that BC5 shifted the GV less in the mutant than in the WT channel (Fig 4e, f), while E219R enhanced BC5 activation (Fig 4e, f). These results support the allosteric pathway coupling Ca^2+^ binding to pore opening (Fig 4a-d) and that BC5 activates the channel by perturbing the pathway. Taken together, our results demonstrate that the molecular interactions at the VSD- CTD interface of BK channels are important in coupling Ca^2+^ binding to pore opening.

## Discussion

We identified BC5 using *in silico* screening in order to modify Ca^2+^ dependent activation of BK channels (Fig 1a, b). Our results showed that BC5 activated the channel in the absence of voltage sensor activation, and not through indirectly modifying voltage dependent gating (Fig 3a, b), but by selectively altering Ca^2+^ dependent activation. BC5 binds at the CTD-VSD interface (Fig 2), away from Ca^2+^ binding sites in the CTD and the pore in the membrane, yet it not only promoted pore opening (Fig 1d) but also interacted with Ca^2+^ binding such that its effect on channel activation was inhibited by Ca^2+^ binding (Fig 3c-f). Thus, BC5 perturbed both the sensor and the pore ends of the Ca^2+^ dependent activation mechanism allosterically by interacting with the pathway that couples Ca^2+^ binding to pore opening. This pathway is built intrinsically in the BK channel protein, since BC5 still activated the Core-MT channel (Fig 3f), in which the entire CTD was removed and replaced with an exogenous short peptide.

We found that BC5 activated BK channels by shifting GV relation (Fig 2f) and increasing open probability at negative voltages (Fig 3b) to a similar extend as 1 µM Ca^2+ 36, 42^. When the dependence of BK channel activation on Ca^2+^ was fit with the MWC model, the Ca^2+^ binding affinity at the open state was ∼1 µM^36, 42^. Thus, BC5 activation of the channel only reached part of the capacity of Ca^2+^ dependent activation (Fig 4d). This is not surprising because Ca^2+^ binding is coupled to pore opening via two major pathways, one is through the CTD-VSD interface, which is affected by BC5, and the other is through the C-linker^21^ (Fig 4a, b). However, in our network and information flow analyses, the highest information flow residues were observed at the VSD- CTD interface (Fig 4 a, b; sFig. 7). These results seem to suggest that the pathway through the VSD-CTD interface may make more contributions to Ca^2+^ dependent activation. However, since the maximal activation by BC5 was less than half of that by Ca^2+^ (Fig 4d), BC5 interaction with the pathway may not be strong enough to even achieve the full capacity of Ca^2+^ dependent activation via the CTD-VSD pathway. This idea is consistent with the result that Ca^2+^ binding inhibited BC5 effects on channel activation (Fig 3c, d), suggesting that the perturbation of the pathway by Ca^2+^ binding was stronger and overrode that by BC5.

Each Slo1 subunit of BK channels contains two Ca^2+^ binding sites in the CTD ^29, 36, 43, 44^, and we found that Ca^2+^ binding to both sites shared the same CTD-VSD allosteric pathway for coupling to the pore. We measured Ca^2+^ effects on BC5 activation of the channel with either Ca^2+^ binding site removed by mutation 5D5N or D367N ^36^. BC5 activated the channel similarly to that in the WT BK in either of the mutant channels in the absence of Ca^2+^, and in 100 µM Ca^2+^ BC5 activation was not abolished as in the WT channel (Fig 3c, d; sFig 8), indicating that the disrupted Ca^2+^ binding to either of the removed site could no longer inhibit BC5 effects via the CTD-VSD pathway. In other words, Ca^2+^ binding to both sites are required to inhibit BC5 effects. These results indicate that both Ca^2+^ binding sites are coupled to the activation gate via the same allosteric pathway that interact with BC5. Since BC5 activated the 5D5N or D367N channels to a similar extent by a similar GV shift (sFig 8), the two Ca^2+^ binding sites may activate the channel with a roughly equal amount via the allosteric pathway that interact with BC5.

BC5 activated BK channels as an allosteric agonist, but it also inhibited the channel as an off-target effect (Fig 1c-e). Opposite to its activation of the channel that was dependent on Ca^2+^ (Fig 3c, d) but not on voltage (Fig 3a, b), the inhibition by BC5 was dependent on voltage but not Ca^2+^ (Fig 1f). While BC5 can be used as an excellent tool to study Ca^2+^ dependent activation mechanisms, it can be also used as an effective BK channel inhibitor in physiological conditions, in which the membrane voltage remains more positive than the K^+^ equilibrium potential so that BK channels are always inhibited. Our results revealed the CTD-VSD interface as an excellent site for allosteric agonists (Fig 2) and the inner pore as a site for inhibitors (sFig 3) of BK channels. These results also suggest that potentially other sites along the allosteric pathways (Fig 4a, b) can be used for allosteric agonists or antagonists to modulate BK channel function. These sites can be targeted for the development of drug therapies for BK channel associated diseases such as neurological disorders associated with either gain-of-function or loss-of-function BK mutations^7^ or the abnormal urinary bladder control^11^.

## Materials and Methods

### Molecular Docking and *in silico* screening

A structure-based *in silico* screening strategy was employed in this study. Specifically, an in-house molecular docking software MDock^45–47^ was used to screen a subset of the Available Chemical Database (ACD, Molecular Design Ltd.) in which each compound has a formal charge of either 1 or 2 (∼ 4×10^4^ compounds), targeting the VSD- CTD interface of the human BK channel (PDB entry: 6v38^20^; Fig. 1a). The Mg^2+^ located at the VSD-CTD interface in the PDB structure was removed during the docking processes. Then, top 2% ranked compounds were manually inspected and 9 compounds were finally selected for experimental tests.

### Mutations and Expression

All of mutations in this study were made by using overlap-extension polymerase chain reaction (PCR) with Pfu polymerase (Stratagene) on the template of the mbr5 splice variant of *mslo1* (Uniprot ID: Q08460)^48^. Sequencing of the PCR-amplified regions is used to confirm the mutation^49^. mRNA was synthesized in vitro from linearized cDNA using T3 polymerase kits (Ambion, Austin, TX). mRNA of mutations was injected into oocytes (stage IV-V) from female *Xenopus laevis* at an amount of 0.05–50 ng/oocyte. The injected oocytes were incubated at 18°C for 2-7 days before electrophysiology recordings.

### Electrophysiology

All experimental data were collected from inside-out patches. The set-up included an Axopatch 200-B patch-clamp amplifier (Molecular Devices, Sunnyvale, CA), ITC-18 interface and Pulse acquisition software (HEKA Electronik, Lambrecht, Germany). Borosilicate pipettes used for inside-out patches were pulled using Sutter P-1000 (Sutter Instrument, Novato, CA) and then wax coated and fire polished with a resistance of 0.5–1.5 MΩ.

The macroscopic currents were collected at the 50 kHz sampling rate (20-μs intervals) with low-pass-filtered at 10 kHz. A P/4 leak subtraction protocol with a holding potential of –120 mV were applied to remove capacitive transients and leak currents (except for experiments in Fig 3b). The solutions used in recording macroscopic ionic currents include: **1)** Pipette solution (in mM): 140 potassium methanesulphonic acid, 20 HEPES, 2 KCl, 2 MgCl2, pH 7.2. **2)** The nominal 0 µM [Ca^2+^]i solution contained about 0.5 nM free [Ca^2+^]i (in mM): 140 Potassium methanesulfonate, 20 HEPES, 2 KCl, 5 EGTA, pH 7.1. **3)** Basal bath (intracellular) solution to make different [Ca^2+^]i (in mM): 140 Potassium methanesulfonate, 20 HEPES, 2 KCl, 1 EGTA, and 22 mg/L 18C6TA, pH 7.2. CaCl2 was then added into the basal solution to obtain the desired free [Ca^2+^]i that was measured by a Ca^2+^-sensitive electrode (Thermo Electron, Beverly, MA).

Gating currents (Fig 3a) were recorded from inside-out patches at the 200 kHz sampling rate and 20 kHz filtration with the same leak subtraction protocol. **4)** Pipette solution for gating currents (in mM): 125 tetraethylammonium (TEA) methanesulfonic, and 2 TEA chloride, 2 MgCl2, 20 HEPES, pH 7.2 and **5)** bath solution for gating currents (in mM): 135 N-methyl-D-glucamine (NMDG) methanesulfonic, 6 NMDG chloride, 20 HEPES, 5 EGTA, pH 7.2.

Single-channel currents (Fig 1g, h) were collected using an Axopatch 200B and sampled at 200 kHz with a Digidata 1322A (Molecular Devices). Filtering for analysis was at 5 kHz^50^.

BC5 (ARG-4-Methoxy-2-Naphthylamine, ordered from Medchemexpress LLC Monmouth Junction, NJ) was dissolved into DMSO to make 100 mM stock solution and then added to recording solutions to the final concentrations. All other chemicals were purchased from Sigma-Aldrich. All the experiments were performed at room temperature (22–24°C).

### Data analysis

Conductance-voltage (GV) relationship was determined by measuring macroscopic tail current amplitudes at –80 mV or -120mV. The GV relationship was normalized to the maximum G value (GMax) and was fitted with the Boltzmann function:

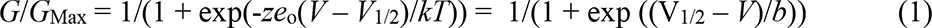

where *z* is the number of equivalent charges that move across the membrane electric field, *eo* is the elementary charge, *V* is membrane potential, while *V*1/2 is the voltage where *G*/*G*Max reaches 0.5, *k* is Boltzmann’s constant, *T* is absolute temperature, and *b* is the slope factor with units of mV. Each GV relationship presented in figures was the averaged results of 3–9 patches and error bars in the figures were the standard error of mean (SEM).

In order to measure limiting slope the open probability (PO) and the total open probability for all the channels N in a patch (NPO) at negative voltages (Fig 3b) were first measured. The currents of all opening events from a patche containing hundreds or thousands of channels during a long pulse (5 or 10s) at each voltage were integrated, and then divided by the single channel current amplitude and total time to obtain NPO.

The estimation of the total number of channels expressed in a membrane is from the equation

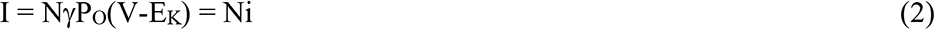

where i represents single channel current and i = PO(V-EK). N was estimated by measuring the macroscopic current at 100 mV in the presence of 100 μM [Ca^2+^]i, where the single channel conductance is 273 pS and PO is about 1.0.

Dose response curves of G-V shifts (Δ*V*1/2) or current inhibition on BC5 concentration (EC50 in Fig 1e, 2e, f, and IC50 in Fig 1e, sFig 3b) was obtained from the fitting to the Hill equation:

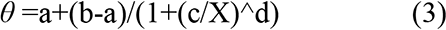

Where *θ* is the expected response at dosage *X*, *a* is the response for dosage at 0, *b* is the maximum response for an infinite dosage, *c*, also known as *EC*50, is the dosage at which the response is 50% of maximum, and *d* is the Hill Coefficient. In all of our final fittings Hill Coefficient was set to 1 since it was close to 1 in the initial fittings.

### Atomistic simulations

The cryo-EM structures of hBK channel in the Ca^2+^-bound (PDB: 6v38) or Ca^2+^-free (PDB: 6v3g) states were used in all simulations reported in this work.^20^ Several short loops absent only in one structure were rebuilt using the other structure as template using the Swiss-PDB server.^51^ For example, A567-E576 and N585-E591 missing in Ca^2+^-bound structure were rebuid using the Ca^2+^-free structure. Missing N- and C-terminal segments as well as several long loops (e.g., V53-G92, A614-V683, and D834-I870) are presumably dynamic and thus not included in the current simulations. Residues before and after the missing segments are capped with either an acetyl group (for N-terminus) or a N-methyl amide (for C-terminus). Standard protonation states under neutral pH were assigned for all titratable residues.

As summarized in Table S1, four sets of atomistic simulations were performed: The Ca^2+^-bound state of BK channel with/without BC5 (*sim 1-3* and *4-6*, respectively) and the Ca^2+^-free state of BK channel with/without BC5 (*sim 7-9* and *10-12*, respectively). The initial binding pose of BC5 was identified with docking study. In simulations *1-3* and *6-9*, a BC5 molecule is placed in binding pocket in each of the four subunits.

All initial structures were first inserted in model POPC lipid bilayers and then solvated in TIP3P water using the CHARMM-GUI web server.^52^ The systems were neutralized and 150 mM KCl added. The final simulation boxes contain about ∼900 lipid molecules and ∼130,000 water molecules and other solutes, with a total of ∼316,000 atoms and dimensions of ∼180 × 180 × 160 Å^3^. The CHARMM36m all-atom force field^53^ and the CHARMM36 lipid force field^54^ were used. The interaction parameters of BC5 were then assigned by identifying similar atom, bond, angle, and dihedral-angle types from similar small molecules within the CHARMM36m all-atom force field^53^. All simulations were performed using CUDA-enabled versions of Gromacs 2018^55, 56^.

Electrostatic interactions were described by using the Particle Mesh Ewald (PME) algorithm^57^ with a cutoff of 12 Å. Van der Waals interactions were cutoff at 12 Å with a smooth switching function starting at 10 Å. Covalent bonds to hydrogen atoms were constrained by the SHAKE algorithm^58^, and the MD time step was set at 2 fs. The temperature was maintained at 298 K using the Nose- Hoover thermostat^59, 60^. The pressure was maintained semi-isotopically at 1 bar at membrane lateral directions using the Parrinello–Rahman barostat algorithm^61^.

All systems were first minimized for 5000 steps using the steepest descent algorithm, followed by a series of equilibration steps where the positions of heavy atoms of the protein, BC5 and lipid were harmonically restrained per CHARMM-GUI Membrane Builder protocol.^52^ Briefly, 6 equilibration step (25 ps for steps 1-3, 100 ps for steps 4-5 and 10 ns for step 6) were performed, where the restrained force constant for proteins and BC5 molecules were set to 10, 5, 2.5, 1.0, 0.5 and 0.1 kcal. mol^-1^.Å^-2^, respectively. For lipids, the phosphorus is restrained with force constants of 2.5, 2.5, 1.0 and 0.5, 0.1 and 0.0 kcal. mol^-1^.Å^-2^, respectively. In the last equilibration step, only protein heavy atoms were harmonically restrained and the system was equilibrated for 10 ns under NPT (constant particle number, pressure and temperature) conditions at 298 K and 1 bar. All production simulations were performed under NPT conditions at 298 K and 1 bar.

### Analysis

Unless stated otherwise, snapshots were extracted every 50 ps for all equilibrium MD trajectories for calculation of statistical distributions. Molecular illustrations were prepared using VMD^62^. Contact probability was calculated with MDAnalysis^63^. Specifically, a residue is considered to contact with BC5 if any of its heavy atom is found within 6 Å of any heavy atom of BC5. The results was first averaged over the four BC5 in four BK subunits, then averaged over three simulations in each condition.

### Dynamical network analysis

The analysis identifies the most probably pathways of dynamic coupling between two selected residues using the Floyd-War-shall algorithm.^38^ For this, each residue (and BC5) is represented as a node of the network. If two residues form a contact (identified with a minimal heavy atom distance cutoff of 5 Å) for greater than 75% of the simulation time, an edge connecting the two corresponding nodes is added to the network. The resulting contact matrix was weighted based on the covariance of dynamic fluctuation (*Cij*) calculated from the same MD trajectory as *wij* = - log(|*Cij*|). The length of a possible pathway *Dij* between distant nodes *i* and *j* is defined as the sum of the edge weights between consecutive nodes along this path. The optimal pathway is identified as the shortest path, thus having the strongest dynamic coupling. Suboptimal paths between *i* and *j* are identified as additional top paths ranked using the path length. The analysis was performed using the Network View^64^. The first 20 optimial and suboptimal paths were selected for analysis presented in this work.

### Information flow

Information flow provides a global assessement of the contributions of all nodes to the dynamic coupling between selected “source” and “sink” nodes.^39, 40^ This analysis complements the dynamic pathway analysis to provide additional insights on how different residues may contribute to sensor-pore coupling. For this, we first construct a network similar to the one described above. Pairwise mutual information was calculated between node *i* and *j* as follows: *M*_*ij*_ = *H*_*i*_ + *H*_*j*_ − *H*_*ij*_. *H*_*i*_ is calculated as 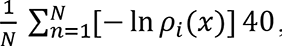, where *ρi(x)* fluctuation density and *x* is the distance to the equilibrium position. Gaussian mixture model (GMM)^65^ is used to estimate the density. The residue network is then defined as *A*ij=*C*ij *M*ij, in which *Cij* is the contact map. To analysis the information flow from source (*S0*) to sink (*SI*) nodes, the network Laplacian, is calculated as *L* = *D* - *A*, where D is diagonal degree matrix: *D*_*ii*_ = Σ_*j*_ *A*_*ij*_.

The information flow through a given node (residue) is defined as 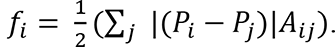 potentials *P* is given by *P* = *L̃*^−1^ *b*. Where *L̃*^−1^ is inverse reduced Laplacian and *b* is b is the supply vector which corresponds to one unit of current entering at the source node that will exit at sink nodes. The magnitude of *fi* thus quantifies the contribution of residue *i* to dynamic coupling between the soure and sink nodes.

## Acknowledgement

This work was supported by grants R01 HL142301 (J.Silva and J.Cui), R35 GM136409 (XZ), R35 GM144045 (J.Chen) and R01 GM114694 (KM) from NIH.

## Author Contributions

G.Z., H.L., J.Shi, M.M., C.A., J.Silva, and J. Cui performed experiments and analyzed data on ion channels expressed in *Xenopus* oocytes. X.X., and X.Z. performed molecular modeling and *in silico* screening. Z.J., and J.Chen performed molecular dynamic (MD) simulations, dynamical network analysis, and information flow. G.Y., and K.M. did experiments and analyzed data on single channel. G.Z., Z.J.,J. Chen, X.Z., X.X, X.Z. and J.Cui. wrote the manuscript.

## Correspondence and request for materials

These should be addressed to J.Cui, X.Z., or J.Chen

---

**Figure S1.**
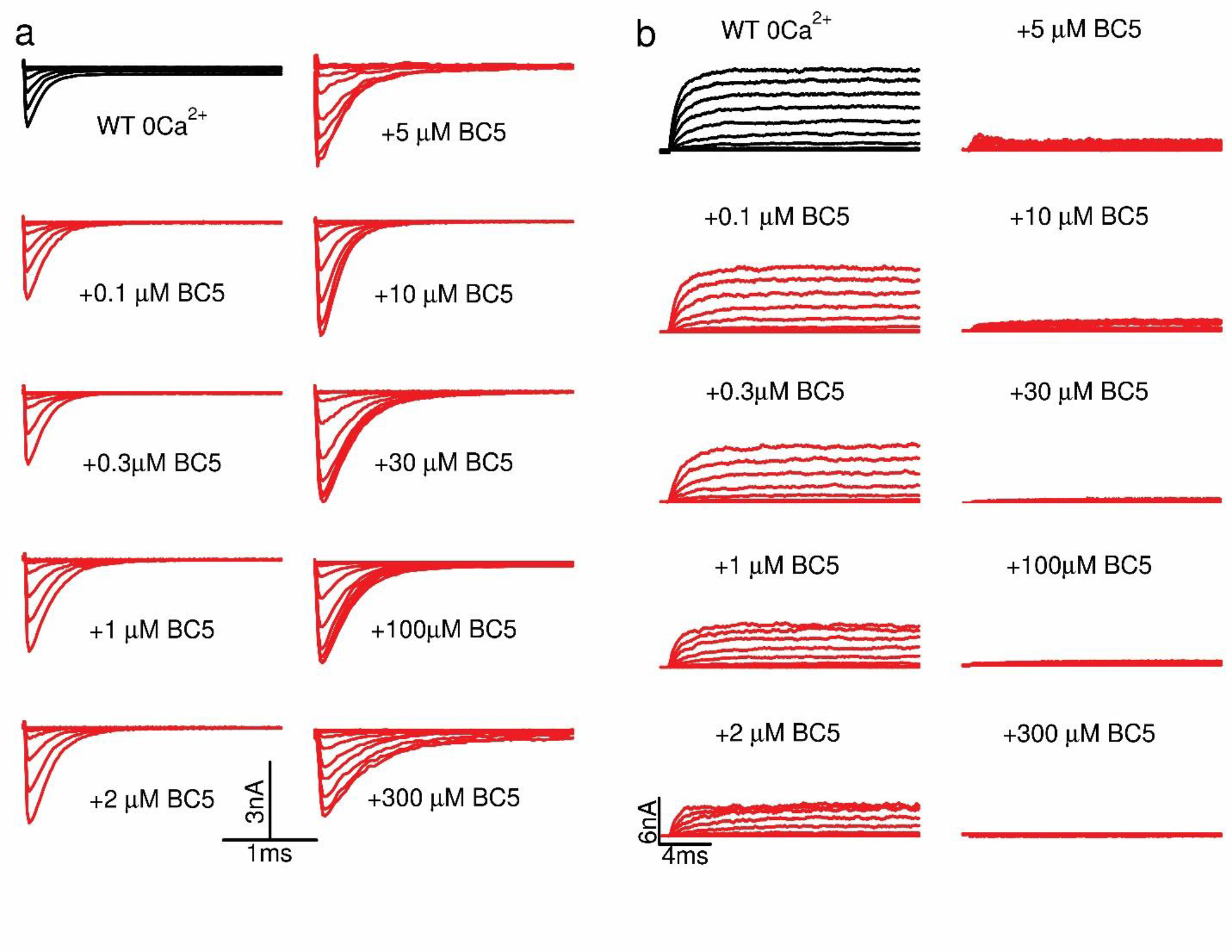
Current traces for BK channels at various BC5 concentrations. **a,** Tail currents. All of tail currents were collected at -80 mV after testing pulses from -80 mV to 200 mV at 20 mV increase. Note that the tail currents started at more negative testing pulses with increasing BC5 concentrations, corresponding to the GV shift in response to BC5 (Fig 1d). **b,** Current traces in response to testing pulses from -80 mV to 200 mV at 20 mV increment. The current amplitudes at +250 mV were used to calculate dose response of BC5 inhibition (Fig 1e).

**Figure S2.**
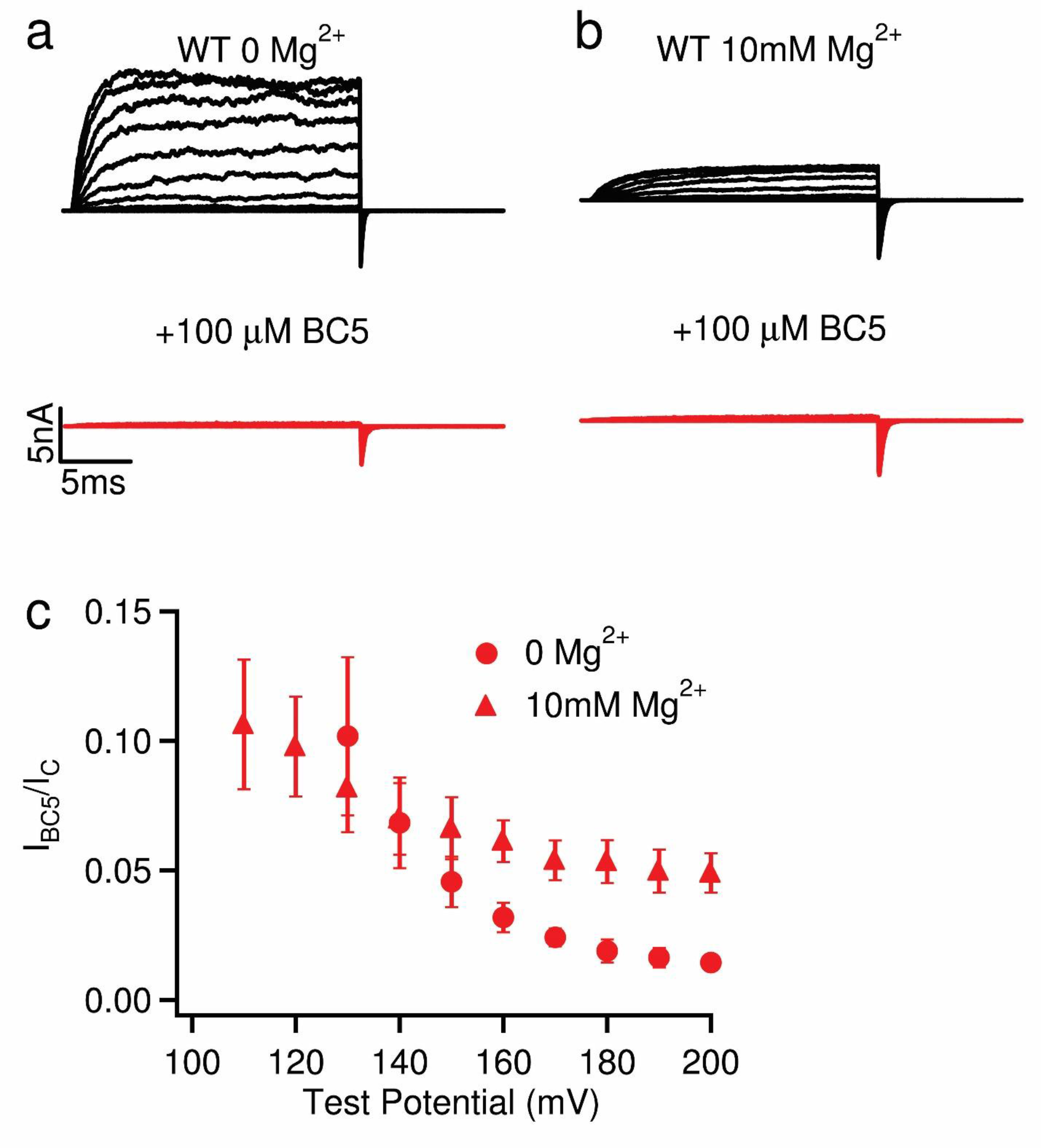
BC5 inhibition of BK channels with and without 10 mM Mg^2+^. **a,** Current traces in 0 Mg^2+^. Voltage pulses: -30mV to 250mV for control (black), -80mV to 200mV in 100 µM BC5 (red), at 20mV increment. The potentials before and after testing pulses were -80 mV. **b,** Current traces in 10 mM Mg^2+^. Voltage pulses were the same as in A except for from -80mV to 200mV. **c,** The ratio of steady state outward current at 100 µM BC5 compared to control at various voltages with (filled triangles) or without (filled circles) the presence of 10 mM Mg^2+^. Note that the BC5 inhibited ≥95% BK currents at 200 mV either with or without 10 mM Mg^2+^. In the presence of Mg^2+^ BC5 inhibition was reduced by a small percentage, possibly due to the interference between Mg^2+^ and BC5, which are both voltage dependent blockers of the channel.

**Figure S3.**
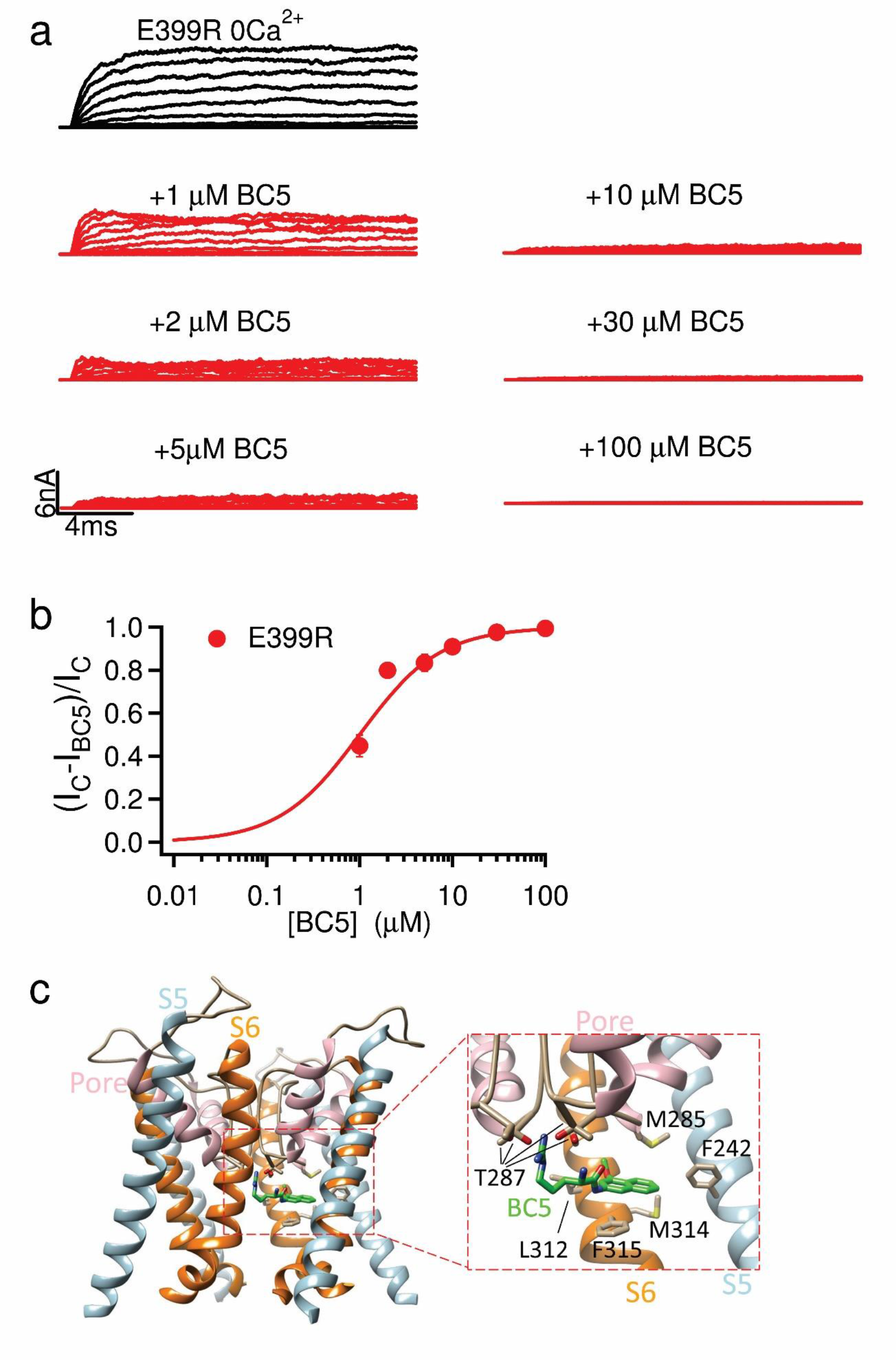
BC5 inhibition of a mutant BK channel with altered BC5 activation, and a possible mechanism of pore blocking. **a,** Current traces of BK channels with the mutation E399R at various BC5 concentrations. **b,** Current inhibition in various BC5 concentrations. Current traces at +200 mV were used to calculate the dose response curve with IC50 of 1.0 µM. Note that the mutation E399R reduced BC5 activation, enhancing the EC50 of V1/2 shift to more than 5 fold (Fig 2f), but the mutation did not enhance IC50 of BC5 inhibition (Fig 1e). **c,** BC5 docked in the pore of the BK channel (PDB 6v3g). S5, the pore helix, and S6 are colored light blue, pink, and orange, respectively. BC5 is represented by the stick model and its carbon atoms are colored green. Residues interacting with BC5 are shown by the stick model and their carbon atoms are colored tan.

**Figure S4.**
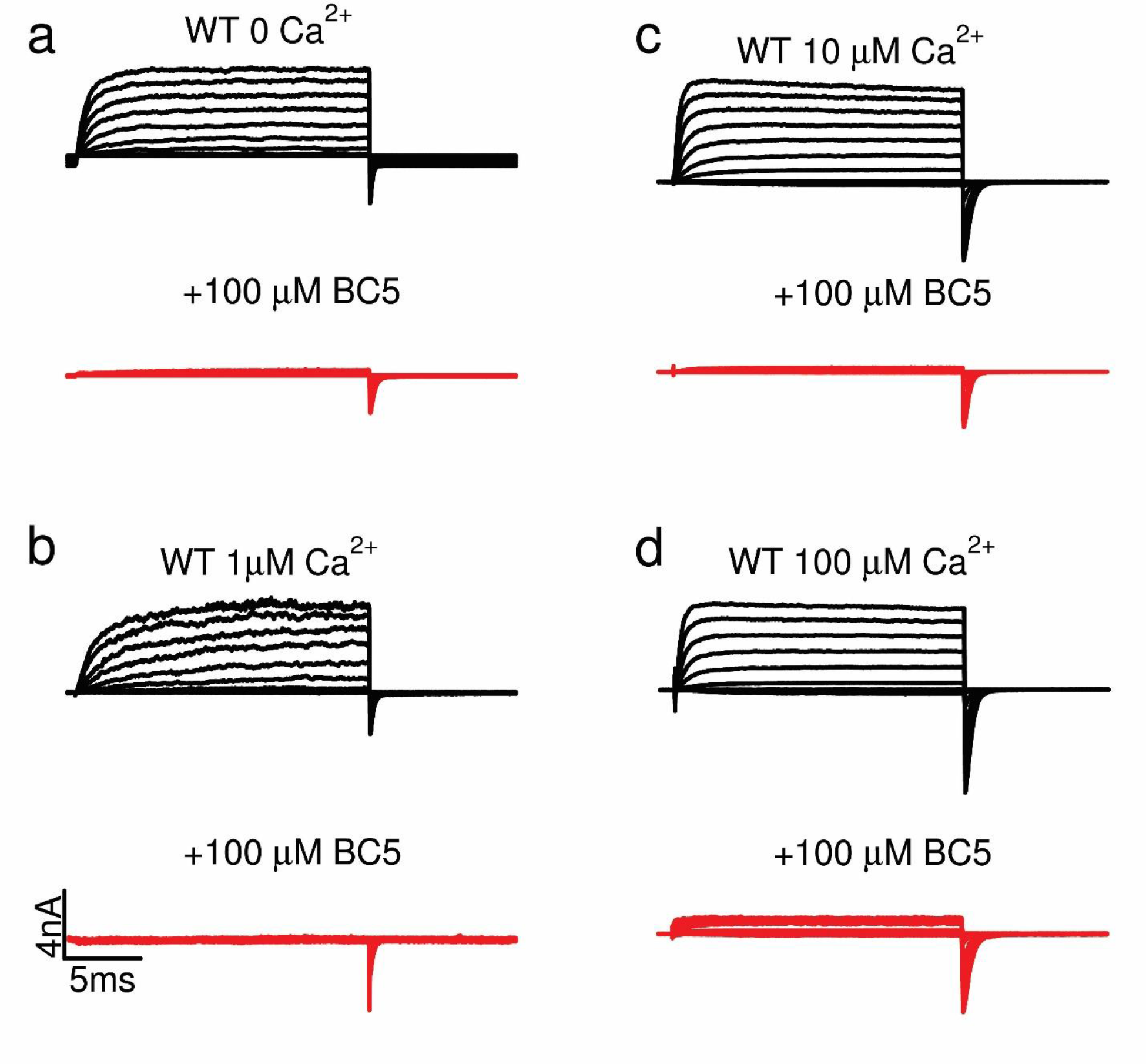
Current traces of BK channels in the presence of different Ca^2+^ concentrations and 100 µM BC5 (corresponding to GV relationships in Fig. 3c).

**Figure S5.**
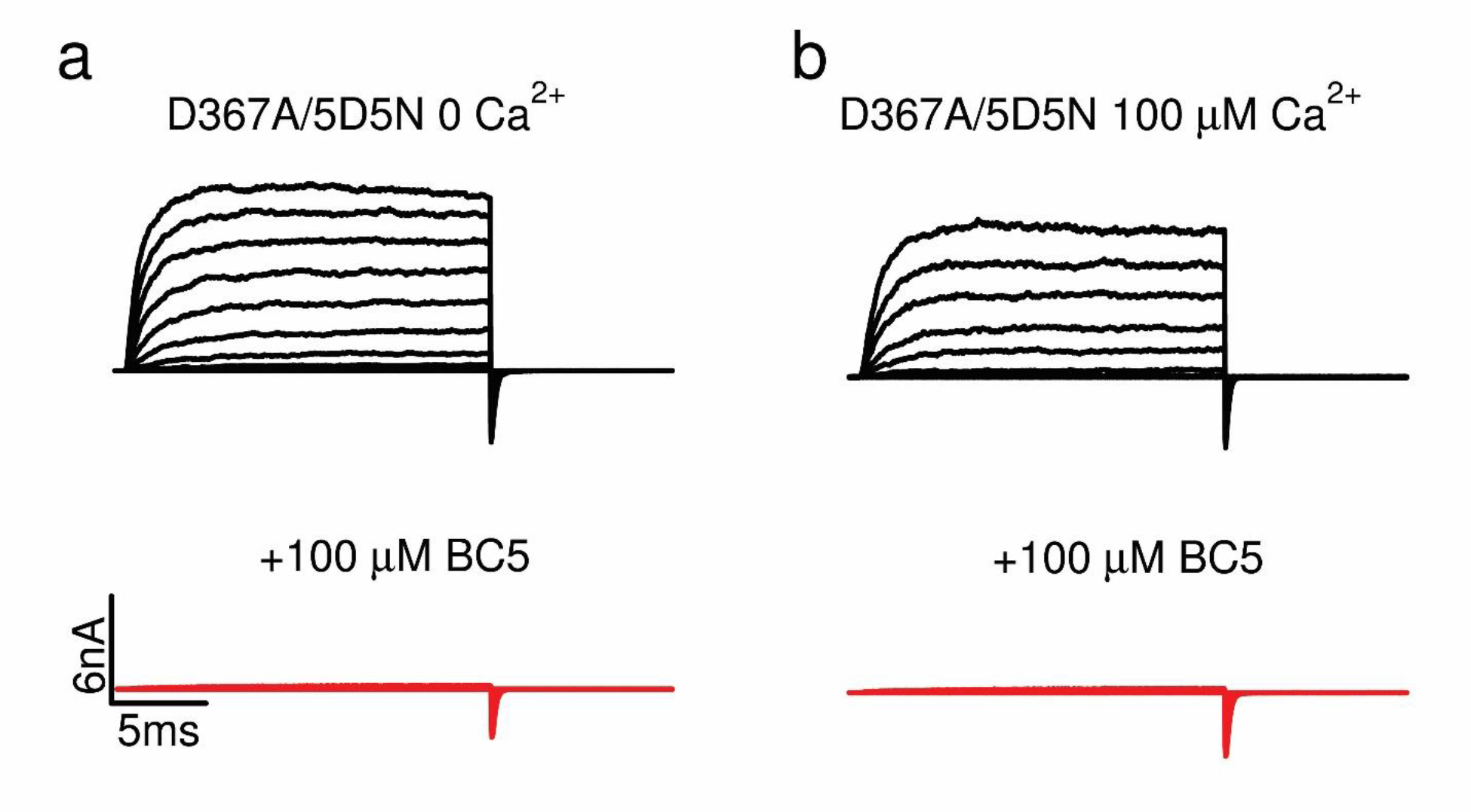
Current traces of mutation D367A/5D5N that removed both Ca^2+^ binding sites of BK channels (corresponding to GV relations in Fig. 3e). **a,** In nominal 0 Ca^2+^, with (red) and without (black) 100 µM BC5. **b,** In100 µM free Ca^2+^.

**Figure S6.**
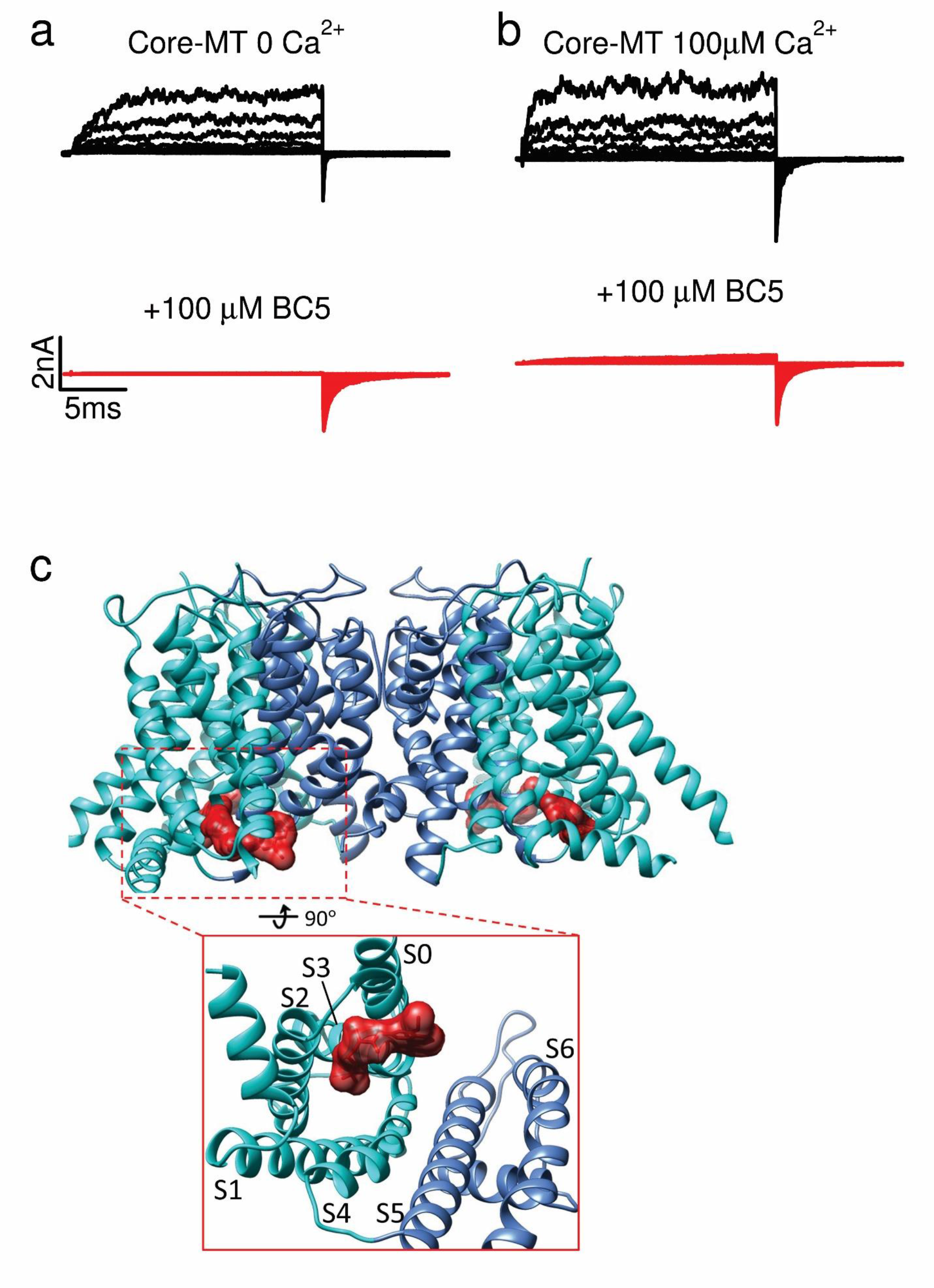
BC5 activation of the Core-MT BK channel. **a, b,** Current traces of Core-MT (corresponding to GV relations in Fig. 3f). **c,** Docking of BC5 on the core-MT (PDB 6v38). BC5 is represented by the surface model and colored red. The tetramer Core-MT is shown by the ribbon model. VSDs are colored cyan and pore-gate domains are colored cornflower blue.

**Figure S7.**
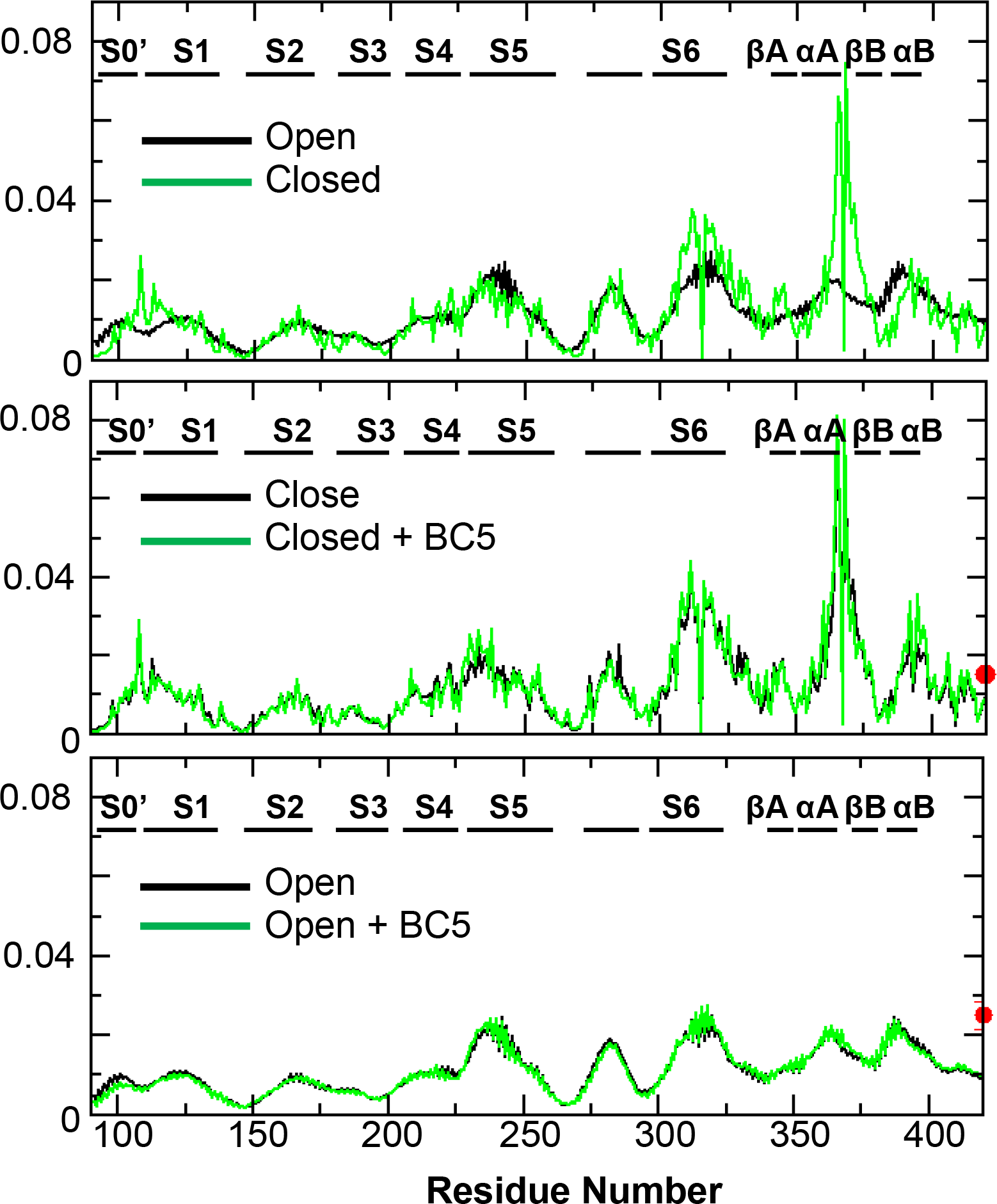
Averaged information flow profiles of BK channels under various conditions. The source and sink nodes are D367 and F315, respectively. Protein specific domains are labeled above the plots. The information though BC5 is also shown as red circle.

**Figure S8.**
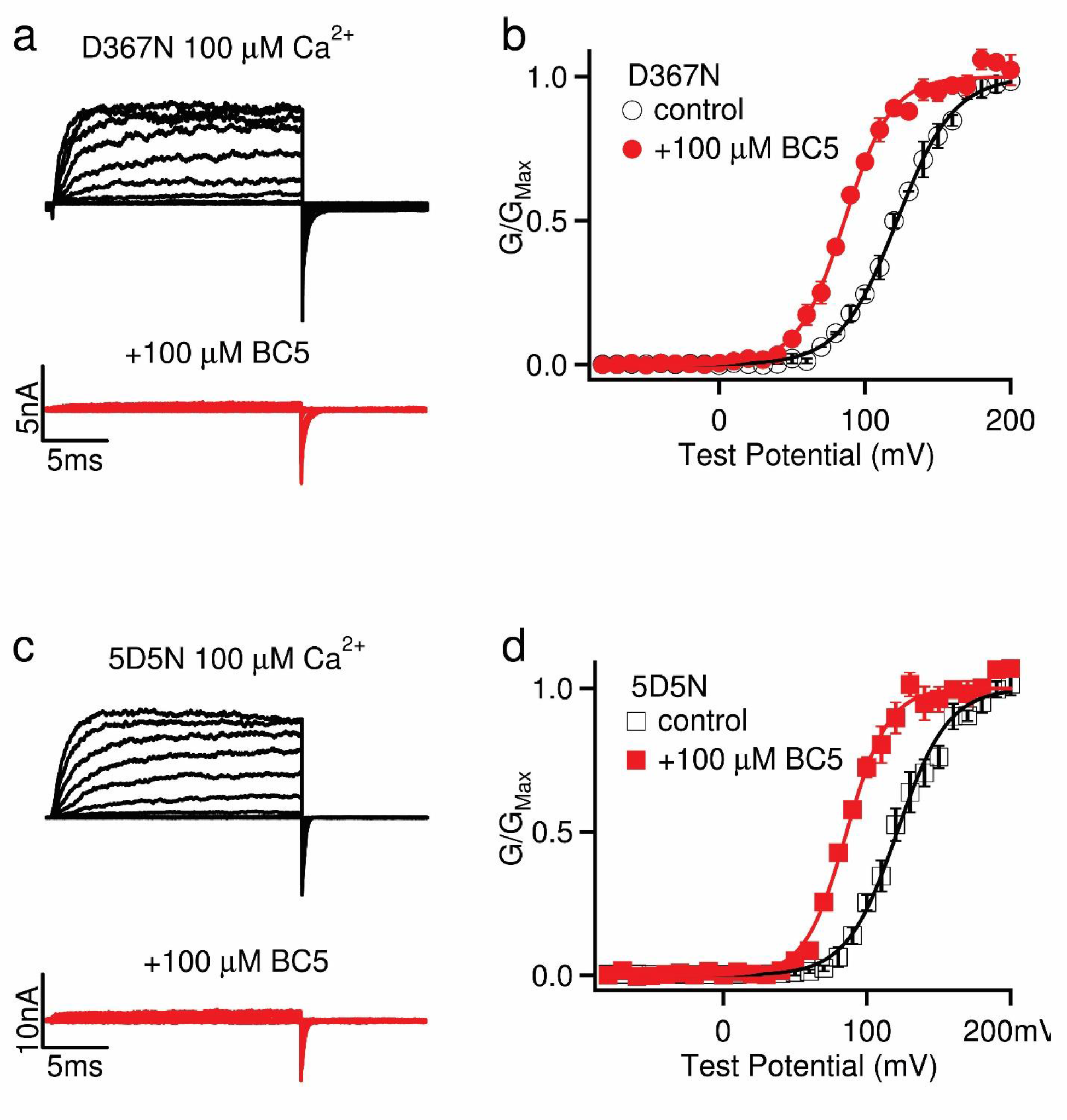
BC5 effect on different Ca^2+^ binding sites. **a,** Current traces at 100 µM [Ca^2+^]i for mutation D367N, ablated Ca^2+^ binding site from RCK1, (control, black) and with BC5 (100 µM, red) at various testing voltages from -80mV to 200mV with 20mV increment. The potential before and after testing pulses was -80 mV. **b,** BC5 effects on the D367N mutant channel in 100 µM [Ca^2+^]i. GV relations were fit with the Boltzmann function (solid lines). V1/2 and slope factor (mV): 122.0 ± 3.2 and 19.5 ± 2.8 for control; and 86.0 ± 3.9 and 16.2 ± 3.4 with 100 µM BC5. **c,** Current traces at 100 µM [Ca^2+^]i for mutation 5D5N, ablated Ca^2+^ binding site from RCK2, (control, black) and with BC5 (100 µM, red) at various testing voltages from -80mV to 200mV with 20mV increment. The potential before and after testing pulses was -80 mV. **b,** BC5 effects on the 5D5N mutant channel in 100 µM [Ca^2+^]i. GV relations were fit with the Boltzmann function (solid lines). V1/2 and slope factor (mV): 126.2± 3.8 and 17.6 ± 2.9 for control; and 83.5 ± 3.2 and 14.5 ± 3.6 with 100 µM BC5.

**Table S1.** Summary of atomistic simulations of BK channels with and without BC5.

| Systems | Ca<sup>2+</sup>/Mg<sup>2+</sup> | BC5 | Initial Structure | Length (ns) |
| --- | --- | --- | --- | --- |
| *sim 1-3* | +/- | + | 6v38 | 200 each |
| *sim 4-6* | +/+ | - | 6v38 | 200 each |
| *sim 7-9* | -/- | + | 6v3g | 200 each |
| *sim 10-12* | -/- | - | 6v3g | 200 each |

**Table S2.** BC5 contact probability of residues in binding pockets

| domain | region | residue | contact probability<sup>1</sup> |  |
| --- | --- | --- | --- | --- |
|  |  |  | Close +BC5 | Open+BC5 |
| VSD | S0' | K 98 | 0.01±0.003 | 0.43±0.04 |
|  |  | G 102 | 0.21±0.09 | 0.44±0.07 |
|  |  | **I 105** | **0.46±0.25** | **0.92±0.09** |
|  | S1 | A 107 | 0.46±0.17 | 0.20±0.01 |
|  |  | L 166 | 0.02±0.01 | 0.52±0.04 |
|  |  | R 167 | 0.29±0.11 | 0.92±0.10 |
|  |  | F 168 | 0.07±0.06 | 0.67±0.12 |
|  |  | A 170 | 0.23±0.11 | 0.95±0.06 |
|  | S3 | A 171 | 0.16±0.06 | 0.95±0.06 |
|  |  | N 172 | 0.22±0.07 | 0.95±0.04 |
|  |  | D 173 | 0.16±0.06 | 0.80±0.17 |
|  |  | F 177 | 0.15±0.10 | 0.87±0.09 |
|  |  | E 180 | 0.28±0.02 | 0.70±0.04 |
|  | S4 | Q 216 | 0.41±0.23 | 0.11±0.05 |
|  |  | E 219 | 0.49±0.21 | 0.27±0.04 |
|  |  | F 223 | 0.42±0.14 | 0.12±0.15 |
| C-linker |  | K 331 | 0.07±0.07 | 0.47±0.08 |
|  |  | G 333 | 0.11±0.12 | 0.42±0.20 |
|  |  | S 335 | 0.09±0.07 | 0.59±0.02 |
|  |  | Y 336 | 0.39±0.13 | 0.78±0.15 |
|  |  | **S 337** | **0.43±0.12** | **0.72±0.09** |
|  |  | V 339 | 0.44±0.14 | 0.39±0.11 |
|  |  | R 342 | 0.30±0.20 | 0.46±0.12 |
| CTD | βB | **E 374** | **0.43±0.16** | **0.71±0.13** |
|  | αB | F 395 | 0.32±0.17 | 0.98±0.01 |
|  |  | **T 396** | **0.42±0.18** | **0.74±0.07** |
|  | βC | Q 397 | 0.29±0.19 | 0.53±0.07 |
|  |  | V 398 | 0.20±0.15 | 0.61±0.15 |
|  |  | **E 399** | **0.41±0.13** | **0.98±0.02** |
|  |  | Y 401 | 0.20±0.08 | 0.87±0.10 |

<sup>1</sup> Only show residues has probability > 40% at east in one state

